# Toxicity by Consortium Revisited: Structural Neighbors and Cross-Family Reach

**DOI:** 10.64898/2026.10.04.756566

**Authors:** Steven M. Muskal, Sanjiv Kumar Jha, Phani Kishore, Prashant Tyagi

**Affiliations:** Eidogen-Sertanty, Inc., Oceanside, CA 92056; AM Green, Plot No. 12, Hitech City, Madhapur, Hyderabad 500081, India; DigitalOcean, Inc., 105 Edgeview Drive, Broomfield, CO 80021; Palo Alto Networks, Inc., 3000 Tannery Way, Santa Clara, CA 95054

**Keywords:** toxicity prediction, structural similarity, nearest neighbors, reverse screening, target breadth, promiscuity, LD50, cytochrome P450, virtual library triage

## Abstract

A compound’s toxicity can be gauged from the toxicities of the compounds it resembles. We showed that in 2003 for rat oral LD50: the compounds in a reference set of 13,645 that most resemble the query form a consortium and vote on its toxicity^1^. This work returns to that method twenty-three years later, on public data, because the reference set used in 2003 is no longer public. Rebuilt on 11,989 curated rat oral LD50 values, the method answers for 67.0% of the held-out compounds carrying an exact value, and 92.1% of those answers land within one log unit; in 2003 the figures were 94% within one log unit on 57.3% of compounds. We then ran it over 28 further endpoints, each recording whether a compound showed a particular toxicity, from cytochrome inhibition to mutagenicity to clinical toxicity: 149,224 measurements on 34,162 compounds. The method answers only when a compound has close enough neighbors, and on those it correctly identifies a median 72.9% of the compounds that showed the toxicity and 96.5% of those that did not. Separately we asked whether a reverse screen carries information about these endpoints beyond molecular size and lipophilicity. We search a compound against a published index of 27,910 ligands solved inside proteins and count the protein families those sites belong to. A compound whose matches reach no protein family inhibits CYP3A4 2.5% of the time; one whose matches span sixteen or more families inhibits 51.2% of the time. Added to heavy atom count, molecular weight and cLogP, the screen raises the mean area under the ROC curve from 0.72 to 0.76 across the 28 endpoints. The OpenADMET consortium has since measured CYP3A4 inhibition on 2,335 compounds, and the same pattern holds. Compounds that resemble known binders of sixteen or more protein families inhibit CYP3A4 24.6% of the time; compounds whose matches reach no protein family inhibit it 7.3% of the time. The more protein families a compound resembles binders of, the more likely it is to inhibit CYP3A4. This association is strong enough to act on before anything is made. Scoring a library on how far its compounds reach across protein families, together with their size and lipophilicity, and setting aside the riskiest fifth removes two thirds of the androgen receptor binders and about half of the CYP3A4 inhibitors. Scoring needs only a two-dimensional structure, with no docking and no conformer generation at query time. Interactive scoring, the data and the pool are at toxpred.ai.

**Graphical abstract:** 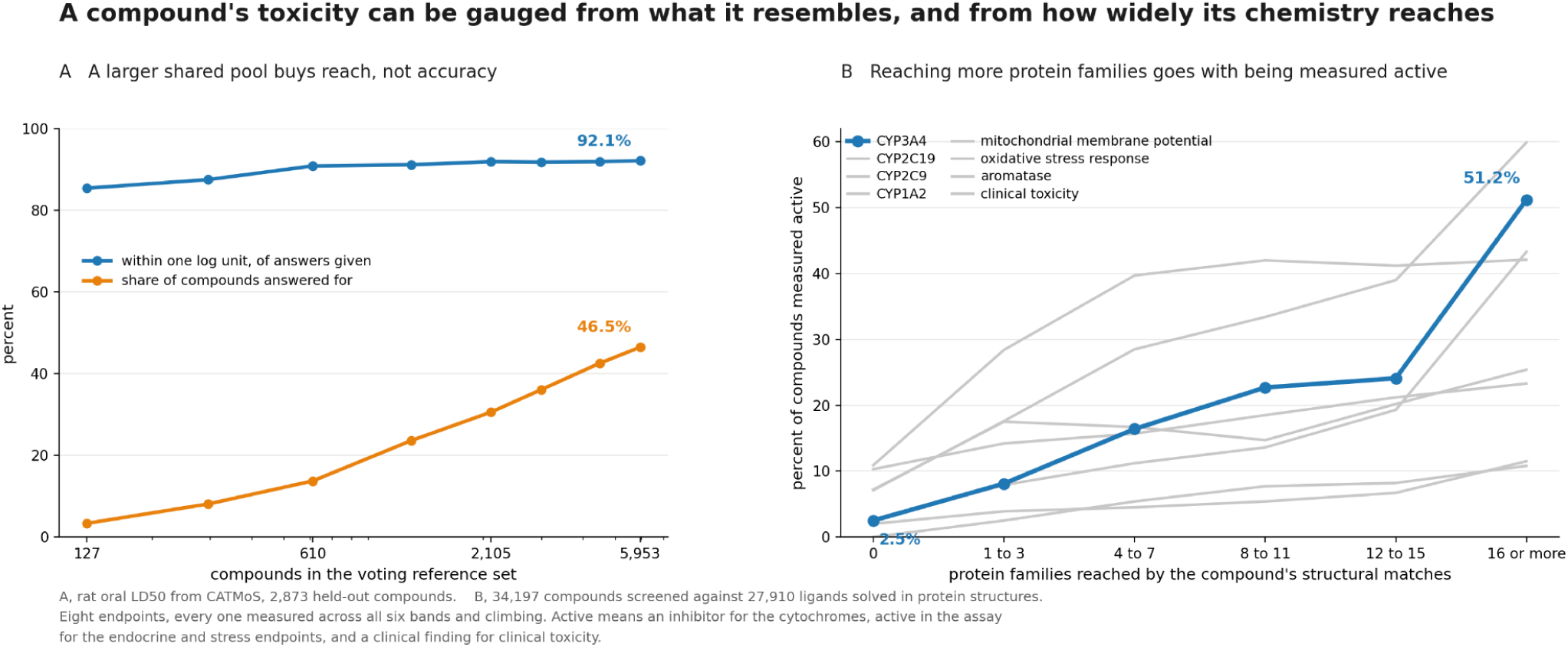

Two signals, both computed from a two dimensional structure. Left, growing the voting reference set from 127 to 5,953 compounds raises the share of compounds answered for from 3.3% to 46.5% while accuracy holds near 92%. Right, the percentage of compounds measured active against the number of protein families their structural matches reach, on eight endpoints where that percentage rises with reach. CYP3A4 runs from 2.5% to 51.2%. Active means an inhibitor for the cytochromes, active in the assay for the endocrine and stress endpoints, and a clinical finding for clinical toxicity.

## 1. Introduction

Safety remains one of the two largest causes of attrition in drug development^2,3^, and the analyses that trace failed candidates back through their histories keep arriving at the same place: compounds fail for reasons that were, in principle, visible early^4^. That observation is what put secondary pharmacology panels into routine early use. A fixed set of receptors, channels, transporters and enzymes is run against a candidate because an undesirable off-target profile can halt development, or lead to market withdrawal when it is found only after approval^5^. Compounds interact with proteins beyond their intended targets, which is why secondary pharmacology panels are used.

What a compound meets beyond its target has been predictable for some time. Chemical similarity alone relates the pharmacology of proteins that share no fold and no family, and about half of the off-target predictions made on that basis are confirmed, in one case explaining a known side effect of a marketed drug through a newly discovered enzyme inhibition^6,7^. Promiscuity itself has been studied directly^8,9^. It is common and it concentrates in certain chemotypes, and how promiscuous a compound appears depends on which activity measurements are admitted and on how the data are selected^8^. Alongside this, a much simpler observation has held up: bulk physical properties predict toxicological outcome. Compounds above a cLogP of 3 and below a polar surface area of 75 square angstroms are more likely to carry an adverse in vivo finding^10^, and lipophilicity drives both promiscuity and attrition generally^11^. Any new signal claiming to predict toxicity has to be shown to add something to those two numbers, and most do not.

The approach we took in 2003 is the opposite of a property rule^1^. It assumes nothing about mechanism and fits nothing. A query compound is fingerprinted, the compounds in a reference set that exceed a similarity cutoff form a voting consortium, and the query inherits the unweighted average of their measured values. When no compound in the reference set is similar enough, the method declines to answer. That design has two consequences that matter more now than they did then. The first is that accuracy and coverage are separate quantities and must be reported together; a method that answers for half a library at 0.82 is not comparable to one that answers for all of it at 0.60. The second is that the reference set is the model. Growing it improves the method with no retraining, so the original paper proposed that organizations pool encoded structure-property pairs without disclosing structures, and we committed to hosting such a pool.

Two things have changed since. The reference set we used, the Registry of Toxic Effects of Chemical Substances^12^, left the public domain and is now licensed commercially. In its place, a curated public collection of rat oral acute toxicity assembled for a modeling collaboration now covers more chemicals^13,14^, and a second resource collects standardized toxicity data over many more endpoints than acute lethality^15^. Both are freely downloadable. The other change is that the crystallographic record has grown to the point where a reverse screen against it is routine^16^. Given a query compound, retrieval over the ligands solved in protein structures returns the proteins those ligands were solved against, and does so in milliseconds, without docking^17^. That gives, for any compound, a computed list of proteins it structurally resembles the ligands of, and therefore a count of how many distinct protein families that list spans.

That count is the quantity this paper is about. Measured promiscuity, read out of an activity database, is contaminated by testing effort: a compound looks promiscuous partly because somebody put it through more panels. A count computed from structure alone has no access to that information. If breadth computed this way tracks toxicity, the correlation cannot be the assay-count artifact, and the argument is one that measured breadth cannot make.

We revisit the original consortium method using public acute toxicity data and 28 additional endpoints. Separately, we examine whether reverse screening provides information about endpoint activity beyond molecular size and lipophilicity, testing it against the possibility that it merely reflects proximity to the crystallographic record. We then combine the retrieval measures with cross-family target preferences and physicochemical descriptors to evaluate their use in virtual library triage. The consortium predictions are evaluated separately and are not inputs to that combined score.

## 2. Methods

### 2.1 The consortium method

The method is unchanged from the original description, to which the reader is referred for its derivation and for the behavior of its parameters^1^. A query compound is compared to every compound in a reference set by Tanimoto similarity over a binary fingerprint. Reference compounds at or above a similarity cutoff form the query’s voting consortium. A prediction is issued only when the consortium has at least a minimum number of members, and is the unweighted mean of their values for a continuous property or the majority of their labels for a categorical one. Weighting votes by similarity gave no gain in the original work and is not used here. Where the method declines to answer, that is recorded rather than filled in, and the share of compounds answered is reported beside every accuracy.

The one substitution is the fingerprint. The 2003 work used 2,048-bit Daylight fingerprints; this work uses Morgan fingerprints of radius 2 folded to 1,024 bits^18^, which are open and standard, scored with the Tanimoto coefficient that remains the method of choice for fingerprint-based similarity^19^.

### 2.2 Toxicity data

Acute toxicity is the curated rat oral collection assembled for the collaborative acute toxicity modeling project and distributed with the OPERA toolset^13,14^. Of its 50,660 records, 11,989 carry experimental data rather than a consensus prediction, and only those are used. 7,948 of them carry an exact point value; the remainder are limit tests or ranges, which cannot enter a regression but determine a hazard category unambiguously. A limit test or a range gives a bound rather than a value, so no error in log units can be computed for it; those records are left out of the regression and its error rather than filled in, and every error reported below is measured against an exact value. Two quantities must be kept apart. Prediction coverage is the share of query compounds for which the method issues a prediction at all; at a required similarity of 0.50 that is 1,764 of the 2,873 held-out compounds, 61.4%. An evaluable prediction is one issued for a compound that also carries an exact value, so that an error can be computed; there are 1,336 of those, 67.0% of the 1,994 compounds carrying an exact value. The series plotted in Figure 1 is the share of evaluable predictions over all 2,873 held-out compounds, and it is labeled as such. 515 of the 11,989 carry no hazard category and 2 could not be fingerprinted, so the sweeps in Figure 1 run on 11,472 compounds, split into a reference set and a held-out set as the legend records. The collection pools readings from different laboratories, strains and protocols. Values are log10 of the LD50 in mg per kg, as the earlier work used, so higher means less toxic. The distribution is close to the one reported in 2003: mean 2.79 against 2.92, standard deviation 0.84 against 0.85.

**Figure 1.**
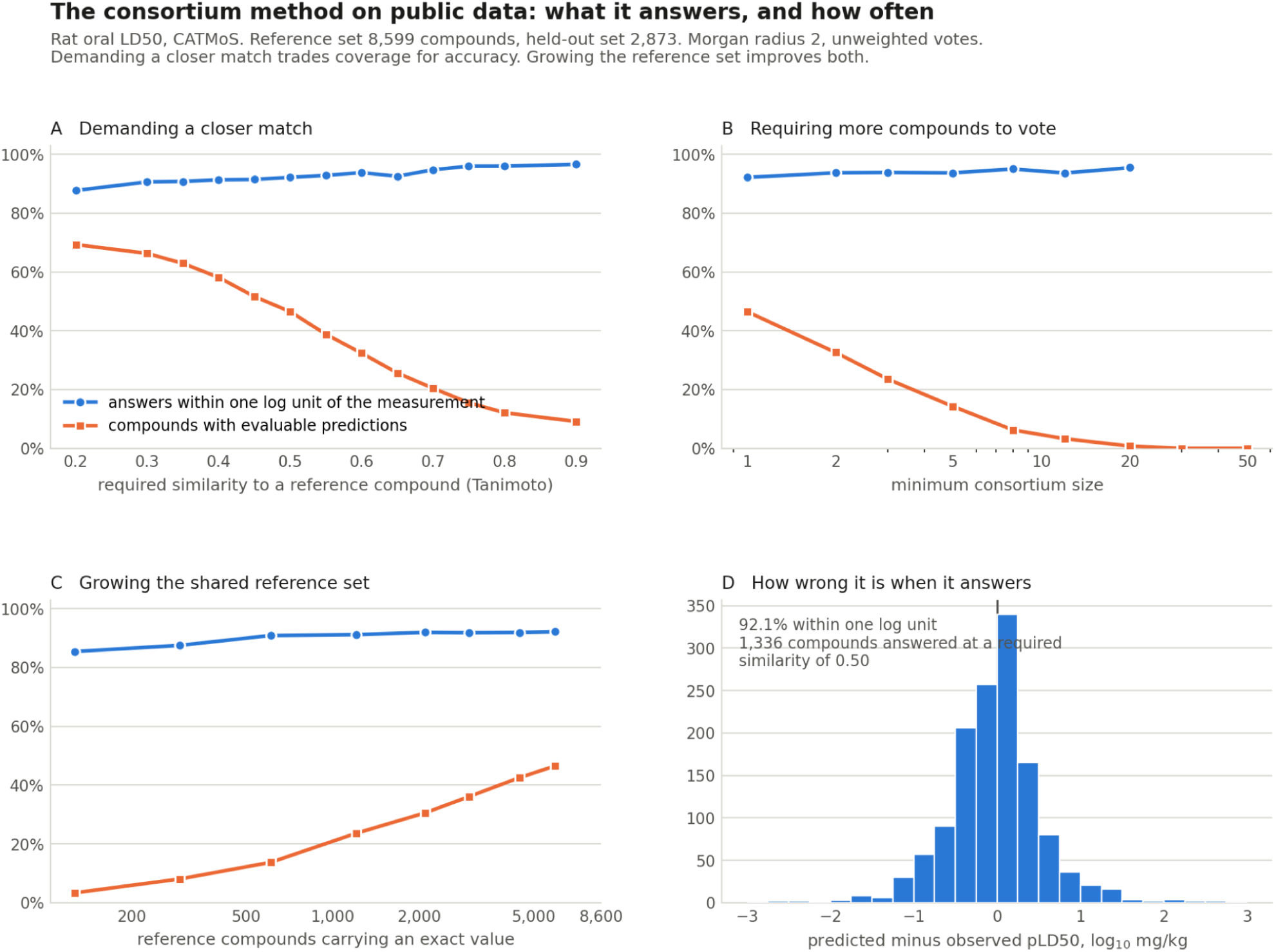
The consortium method on public data. Rat oral LD50 from CATMoS, reference set 8,599 compounds of which 5,953 carry an exact LD50 and may vote, held-out set 2,873, Morgan radius 2 at 1,024 bits, unweighted votes. In every panel the blue series is the share of answers landing within one log unit of the measurement and the orange series is the share of all 2,873 held-out compounds with an evaluable prediction, meaning the method answered and the compound carries an exact value. Prediction coverage, which does not require an exact value, is higher; at a required similarity of 0.50 it is 61.4%. **A**, raising the similarity a reference compound must meet before it may vote. **B**, raising the number of compounds that must vote before an answer is issued. **C**, growing the voting reference set from 127 to 5,953 compounds at a fixed required similarity of 0.50; accuracy rises from 85.4% to 90.9% by 610 compounds and then holds, while the share answered rises the whole way from 3.3% to 46.5%, which is the property a shared pool is built on. Coverage is expressed over all 2,873 held-out compounds, including the 879 whose measurement is a limit test or a range and carries no exact value. **D**, the distribution of prediction error at that operating point, 92.1% of answers within one log unit.

The further endpoints are 30 of the curated endpoint datasets distributed for download by TOXRIC^15^, covering cytochrome P450 inhibition for five isoforms, cardiotoxicity at four thresholds, mutagenicity, carcinogenicity, hepatotoxicity, twelve nuclear receptor and stress response assays, eye irritation and corrosion, respiratory, developmental, reproductive and clinical toxicity. The release carries 156,460 records before duplicate resolution. Resolving duplicates to one measurement per compound and endpoint, and excluding one record that carries no InChIKey, the 30 sets hold 149,557 measurements over 34,197 compounds. 28 of the 30 carry enough compounds to be scored; those 28 hold 149,224 measurements over 34,162 compounds, and every count reported below is drawn from them. The two set aside are developmental toxicity, with 187 measurements, and reproductive toxicity, with 146. The cytochrome inhibition endpoints are derived from IC50 measurements pooled across laboratories, and such collections have been shown to carry uncertainty of the order of a log unit^20,21^. Splits are drawn at the compound level.

### 2.3 Reverse screening and target breadth

Structural matches are retrieved from a published index of 27,910 ligands solved in protein structures, each carrying the UniProt accessions of the proteins it was solved against and spanning 28,632 distinct proteins^17^. The Protein Data Bank^16^ is updated weekly on Wednesdays, and the index is rebuilt shortly thereafter; every retrieval reported here, including the external set of section 3.9, used the release of 20 September 2026. The query and every indexed ligand are represented by a pharmacophore fingerprint predicted from two-dimensional structure, so that no conformer ensemble is generated at query time. Every indexed ligand at or above a Tanimoto similarity of 0.50 is kept, with no limit on how many a query may match, which is what the served implementation does. Families reached is the number of distinct level-2 protein families among the proteins those matches carry, using the family assignments of the released cross-family comparator^22^. A target annotated with two families counts as two, so combined labels are split before counting. A matched protein contributes only if it carries a level-2 assignment, so a compound can hold matches and still reach no family; reaching no family means no matched protein carried an assignment, not that nothing matched. The index spans 32 families and the most any compound reaches is 29.

Two additional screens test whether the count reflects a compound’s own chemical series. In the first, every indexed ligand sharing the query’s Bemis-Murcko scaffold^23^ is removed before retrieval. In the second, every indexed ligand at or above 0.70 similarity to the query is removed before retrieval. Both are computed for every compound.

### 2.4 Cross-family preference

The second signal is taken from a released cross-family target comparator^22^, which given one compound and two proteins returns which of the two the compound prefers. A panel of 19 safety-relevant proteins spanning distinct families was assembled, every pair scored for every compound, and each protein’s win rate standardized across compounds, so that what remains is how unusually a compound is preferred at that protein relative to every other compound in the set rather than a property of the protein itself.

Two phrases used later need defining here. A compound ranks highly at a panel protein when its standardized win rate there is more than one standard deviation above the mean across compounds. A panel protein is confirmed by both when the retrieval reaches it and the comparator also ranks the compound highly at it, and families confirmed by both counts the distinct families among those proteins. The comparator is used exactly as released and is not refitted on any compound here.

### 2.5 Evaluation and what enters each reported score

Comparisons against molecular size and lipophilicity use heavy atom count, molecular weight and cLogP on the identical compounds. Enrichment is the share of toxic compounds among those flagged, divided by the share in the library they came from.

Three different scores are reported and they do not share inputs. The consortium prediction of section 2.1 is computed from the reference set alone and has no fitted step. The marginal-value models of section 3.3 are random forests of 200 trees with a minimum of three samples per leaf, fitted for each endpoint separately, and they are scored by the area under the ROC curve. The combined score of section 3.7 is a gradient boosting classifier of 200 stages at depth three, also fitted for each endpoint separately.

The combined score takes fifteen inputs. Three describe size and lipophilicity: heavy atom count, cLogP and molecular weight. Five come from the reverse screen: families reached, distinct proteins reached, how many of the 47-protein antitarget list those proteins include, the number of indexed ligands matched, and the best similarity among those matches. Seven involve the cross-family comparator of section 2.4: how many panel proteins the compound ranks highly at, the largest standardized win rate, the spread of those win rates, how many panel proteins are reached, how many are confirmed by the retrieval and the comparator together, how many families are confirmed by both, and the compound’s mean rank over the proteins it reaches. Consortium predictions are not among the inputs, so no result in section 3.7 depends on them.

The combined score is fitted on 70% of each endpoint’s compounds and reported only on the 30% held back, drawn once at the compound level with a fixed seed, and an endpoint enters that analysis only if it carries at least 800 compounds with at least 60 in each class, which is why fewer than 28 endpoints appear in some panels. The marginal-value models are fitted on half of each endpoint’s compounds and reported on the other half, drawn the same way, and an endpoint enters those analyses if it carries at least 500 compounds with at least 25 in each class. Where the consortium declines to answer, that compound is recorded as unanswered and is never imputed.

## 3. Results

### 3.1 The method behaves as it did in 2003

Figure 1 repeats the three parameter sweeps of the original paper on the public reference set. Raising the similarity a reference compound must meet before it may vote raises accuracy from 87.7% to 96.6% of answers within one log unit, while the share with an evaluable prediction falls from 69.3% to 9.2%. Raising the number of compounds that must vote does the same thing more sharply, carrying the predictive correlation from 0.57 with a single voter to 0.88 with twenty while coverage all but disappears. Both reproduce the shape of the original figures.

The third sweep is the one the pooling argument rests on. Growing the reference set from 127 to 5,953 compounds carrying a measured LD50 raises accuracy from 85.4% to 90.9% by 610 compounds, after which it holds between 91.2% and 92.1%, while the share with an evaluable prediction rises the whole way, from 3.3% to 46.5%. Past the first few hundred compounds a larger shared reference set buys reach rather than accuracy, so contributing to a common pool is worth something to every contributor. At a required similarity of 0.50 the method answers for 1,336 of the 1,994 held-out compounds carrying an exact value, 67.0% of them, and 92.1% of those answers land within one log unit of the measurement. In 2003 the figures were 94% within one log unit, on the 57.3% of compounds the method would answer for.

### 3.2 Twenty-eight further endpoints

Nothing about the method is specific to lethality. Figure 2 applies it to the 28 endpoints, with no setting retuned except the required similarity, which every endpoint sets by the same rule. The same trade appears in every one: tightening the required match raises the share called correctly and lowers the share answered. Across the 28, the median share called correctly rises from 86.0% to 96.6% as the required match tightens, with most of that rise spent by 0.60, while the median share answered falls to 8.6% and is still falling at the tightest match tested.

**Figure 2.**
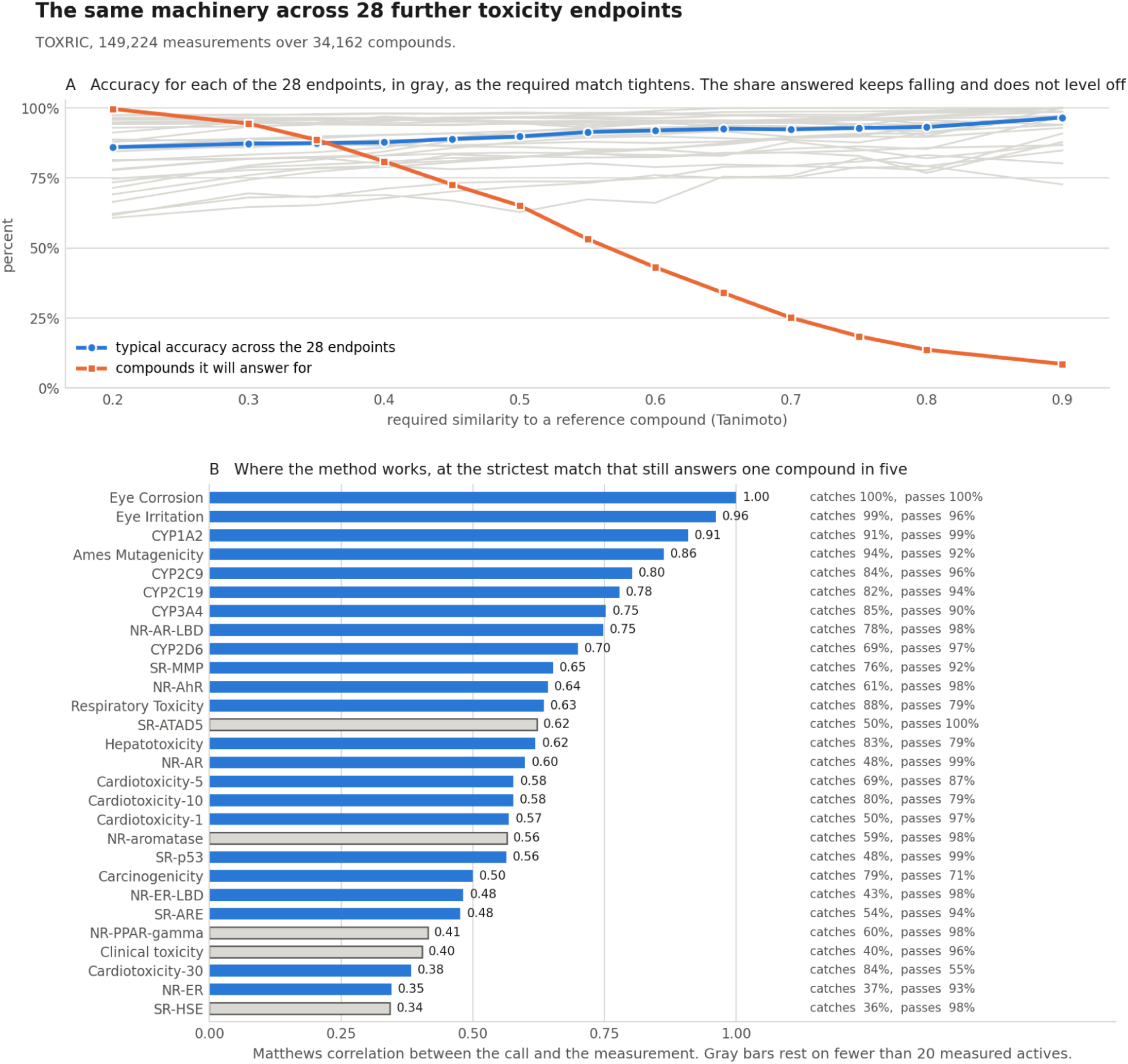
The same machinery over 28 further endpoints from TOXRIC, 149,224 measurements over 34,162 compounds. No setting is retuned except the required similarity, which every endpoint sets by the same rule. **A**, one gray line per endpoint giving the share of its answers called correctly as the required similarity tightens, with the median across the 28 in blue and the median share of compounds answered for in orange. Accuracy gains are mostly spent by a required similarity of about 0.6, while the share answered keeps falling to 8.6% at 0.90 without leveling off, which is why the axis runs to zero. **B**, the 28 endpoints ranked by the Matthews correlation between the call and the measurement, at the strictest match that still answers one compound in five, with the share of measured actives caught and the share of measured inactives passed printed beside each. Matthews correlation is used rather than percent correct because the labels are lopsided, 87 to 13 on CYP2D6, and that ratio records which compounds were tested and reported rather than the molecule; it draws on all four cells of the table, so no answering strategy inflates it. Gray bars rest on fewer than 20 measured actives and are too thin to rank.

These endpoints carry lopsided labels. 87% of the CYP2D6 compounds share one label, which reflects which compounds were tested and reported rather than any property of chemistry, so the share called correctly on such data records that lopsidedness more than it records the molecule. Figure 2B therefore gives the two rates that are unaffected by it: of the compounds measured active, the share the method calls active, and of those measured inactive, the share it calls inactive. Among the compounds answered for, at the strictest match that still answers one compound in five, eye corrosion, Ames mutagenicity and CYP1A2 inhibition each catch more than nine in ten of the measured actives while passing a comparable share of the inactives; CYP3A4 and hepatotoxicity fall to between eight and nine in ten. Figure 2B prints both rates beside every endpoint, and Table S1 gives the required similarity, the share answered and both rates for all 28.

Figure 2B ranks the 28 by Matthews correlation, from 1.00 on eye corrosion down to 0.35 on the estrogen receptor assay among the endpoints carrying at least 20 measured actives. The two rates beside each bar say which way an endpoint fails. Where the method is weak, it is weak in one direction: on the estrogen receptor assay and the heat shock response it calls roughly a third of the measured actives active while passing more than nine in ten of the inactives. It misses actives rather than raising false alarms. Cardiotoxicity at 30 micromolar is the one endpoint that fails the other way, catching most of the actives while passing barely half the inactives. Five endpoints carry fewer than 20 measured actives at this operating point and are drawn gray; clinical toxicity rests on five, so its rates are read as counts rather than ranked.

### 3.3 Reaching more protein families

Figure 3 groups every compound by how many distinct protein families its structural matches reach, and reads off the percentage of each group measured active. For CYP3A4 inhibition the rate climbs from 2.5% among compounds reaching no family to 51.2% among those reaching sixteen or more, a twentyfold difference across six bands. CYP2C9, CYP2C19 and clinical toxicity climb the same way, and panel A plots them with CYP1A2 and cardiotoxicity at 5 micromolar; Table S2 gives every band for all 28 endpoints.

**Figure 3.**
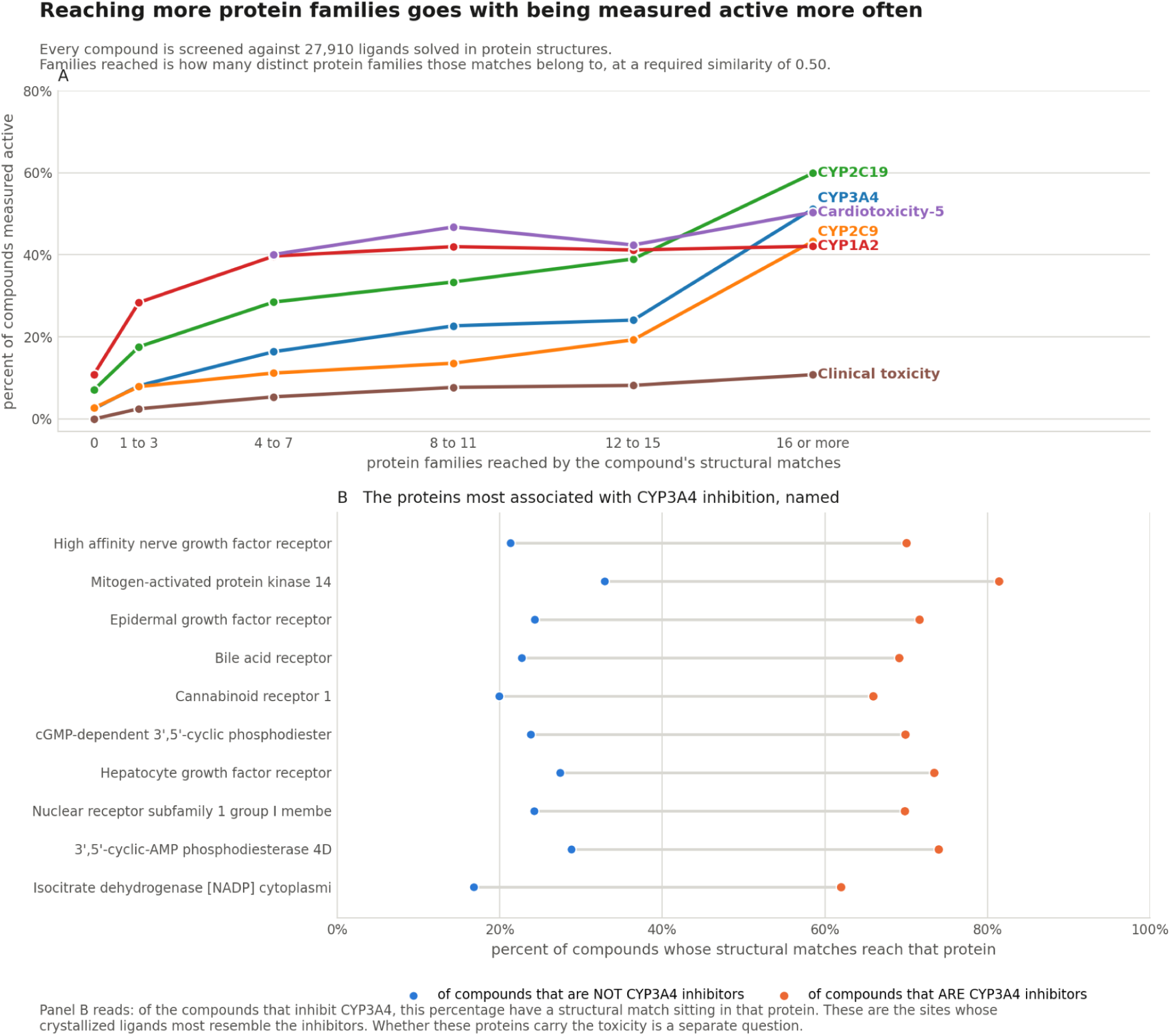
Reaching more protein families goes with being measured active more often. Every compound is screened against 27,910 ligands solved in protein structures, and families reached counts the distinct protein families those matches belong to at a required similarity of 0.50. **A**, percent of compounds measured active in each band of families reached, for six endpoints where the rate climbs: CYP1A2, CYP2C9, CYP2C19, CYP3A4, cardiotoxicity at 5 micromolar and clinical toxicity. **B**, for CYP3A4, the proteins whose crystallized ligands most resemble the inhibitors, with the percent of inhibitors and of non-inhibitors whose matches reach each one. Active means an inhibitor for the cytochromes, active in the assay for the endocrine and stress endpoints, and a clinical finding for clinical toxicity. The endpoints that end no higher than they start are given band by band in Table S2.

Six endpoints do not follow it. Eye corrosion, eye irritation, respiratory toxicity, Ames mutagenicity, carcinogenicity and hepatotoxicity end no higher than they start, and CYP2D6 takes a third shape, rising to a peak at four to seven families and easing back. CYP1A2 rises but stops early, reaching 39.7% by four to seven families and holding near 42% above that. The common thread is molecular size. Compounds reaching no family are small, a median of nine to fifteen heavy atoms, while those reaching sixteen or more are drug sized, a median of twenty-four to thirty-one. The direction of the association differs by endpoint. Compounds in the lowest bands are generally smaller, which may contribute to that difference. These analyses do not establish the mechanisms responsible for the reversed trends.

The count is not standing in for size. Adding the reverse screen to a model already holding heavy atom count, molecular weight and cLogP raises the mean area under the ROC curve from 0.72 to 0.76 across the 28 endpoints, a gain on every one of them; the largest gains fall on clinical toxicity, PPAR gamma binding and cardiotoxicity at 1 micromolar. The reverse screen is five quantities, of which families reached is one, and they are not interchangeable. Added to size on its own, distinct proteins reached is the strongest single column and families reached is weaker; once the other four are in the model, families reached adds almost nothing. Family count is the interpretable summary of the screen, not the quantity carrying it.

Reaching more families does not mean more kinds of toxicity. This was measured on the 5,882 compounds carrying ten or more measured endpoints, so that the comparison is like for like. Across the six bands the share of a compound’s endpoints that come back toxic runs between 8.4% and 12.6%, with no consistent direction; the rank correlation against families reached is 0.05. The number of categories a compound is positive in does not rise either. It runs from 0.95 in the lowest band to 0.81 in the highest, against 2.40 and 2.01 categories measured, at a rank correlation of -0.04. The signal is specific to particular endpoints, not a general tendency to be toxic.

Restricting the count to families a safety pharmacology panel exists to cover, the cytochromes, nuclear receptors, the Family A G protein-coupled receptors, the three ion channel families and the two transporter families, does not improve it either. Counting only those, and rebanding because the panel spans too few families to reach sixteen, CYP3A4 and CYP2C9 climb over almost the same range as they do over all families. Breadth itself carries the signal, whichever families are reached.

Figure 3B names the proteins. Among compounds that inhibit CYP3A4, a structural match sitting in the high affinity nerve growth factor receptor, mitogen-activated protein kinase 14 or the epidermal growth factor receptor is between two and a half and three times as common as it is among compounds that do not, and the panel prints both rates. For CYP2D6 the leading proteins are different, headed by dipeptidyl peptidase 4, son of sevenless homolog 1 and the extracellular calcium-sensing receptor. These are the sites whose crystallized ligands most resemble the inhibitors. Twelve compounds from the top band are drawn in Figure S1.

### 3.4 The trend is not a compound finding its own analogs

A compound that reaches many protein families might simply be a close analog of well-studied ligands, and the crystallographic record is composed of exactly such ligands. Figure S2 tests that possibility directly. Each compound is screened three times: once with the index as it stands, once with every indexed ligand sharing its Bemis-Murcko scaffold deleted first, and once with every indexed ligand above 0.70 similarity to it deleted first.

Scaffold and analog exclusions produce little change in the endpoint rates. No CYP3A4 band shifts by more than 0.4 of a percentage point under either blinding, and across all 28 endpoints the largest shift in any band is 3.2 points. The test is not a weak one: 35.2% of the compounds share a scaffold with an indexed ligand and 47.0% have a match at or above 0.70 similarity, so a great deal was available to remove. Removing it changed families reached for 4.1% and 6.4% of compounds respectively, by 0.06 and 0.10 families on average. Breadth is not carried by a compound’s nearest relative; a compound reaching sixteen families reaches them through many different chemotypes, and deleting its own series barely alters the count.

### 3.5 Robustness to index subsampling

The reference set is the model, and growing it buys reach. The crystallographic index is the second pool this work draws on, and the same question can be asked of it. Subsampling the published index uniformly at random to three quarters, a half and a quarter of its 27,910 ligands, and rescreening every compound at each size, leaves the separation where it was; averaged over the 28 endpoints, and from three independent draws at each reduced size, the area under the curve changes by less than 0.01 across the whole range. Families reached is binned to six quantile bins at every size, because a smaller index gives the count fewer distinct values and ties depress a rank statistic; unbinned the answer is the same. While compounds do reach fewer families as the index shrinks, from 9.9 on average to 6.5, they are ranked no less well for it.

Within the present composition of the index, the ranking is therefore robust to how much of it is retained. This tests redundancy among the ligands now in the record. It does not establish how the count behaves on protein families the record does not yet contain, and a release that adds new families rather than more of the same chemistry is the case this experiment cannot speak to.

### 3.6 Complementary information from retrieval and target preference

Structural reach and the comparator’s preference could be two views of one thing, and they are not. Among compounds whose matches reach few families, how many safety panel proteins the comparator ranks them highly at barely moves the CYP3A4 rate. Among compounds reaching many families the same question nearly doubles it, from 33.1% to 61.3%. The comparator separates compounds only among those whose matches already reach many families. CYP2C9 behaves the same way and CYP2C19 does not.

Counting proteins confirmed by both signals rather than by either alone beats both on every cardiotoxicity threshold, which is the group of endpoints whose panel proteins include the cardiac ion channels, and loses to structural reach alone on four of the five cytochromes. Both signals therefore enter the combined score used below.

### 3.7 Running a library through the score

Figure 4 puts the score in the terms a virtual library triage would use. Out of every 100 compounds, the tenth the score flags holds six times as many androgen receptor actives as the library it came from, three times as many CYP2D6 inhibitors, and close to three times as many clinically toxic compounds; panel A prints the library rate, the flagged rate and the multiplier for every endpoint. That flagged tenth carries 60% of all the androgen receptor ligands in the set and 33% of the CYP2D6 inhibitors.

**Figure 4.**
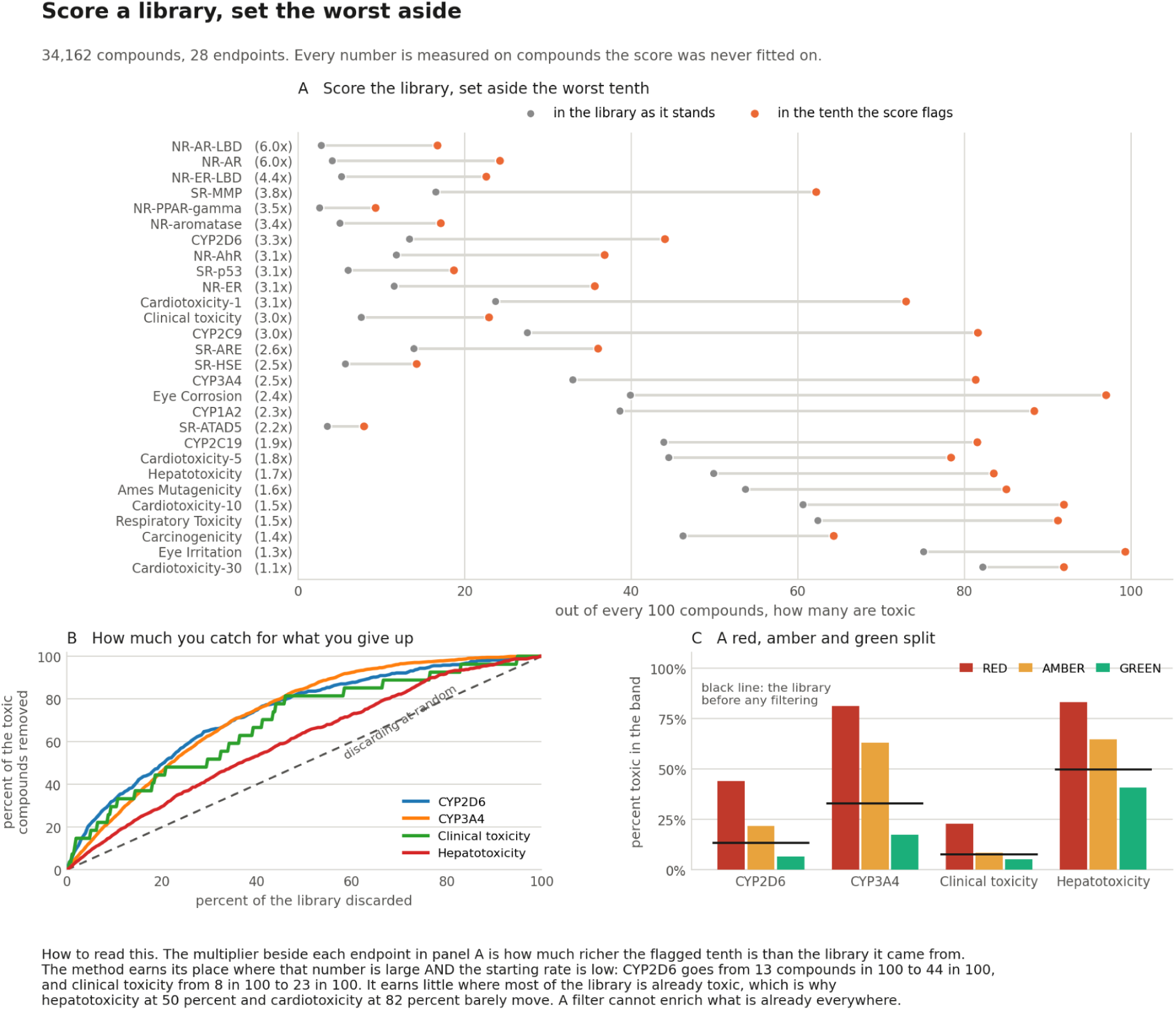
**Running** a library through the score, measured on compounds it was never fitted on. **A**, out of every 100 compounds, how many are toxic in the library as it stands and how many are toxic in the tenth the score flags, with the multiplier beside each endpoint. **B**, discarding the worst scoring compounds, the share of the toxic ones removed against the share of the library given up, with the diagonal marking what discarding at random would achieve. **C**, percent toxic in a red, amber and green split at the worst scoring tenth, the next fifth, and the remainder, with the black line marking the library before any filtering.

Discarding the worst scoring fifth removes two thirds of the androgen receptor actives and about half of the CYP2D6 and CYP3A4 inhibitors, against a fifth for discarding at random. Splitting the library into red, amber and green bands at the worst tenth, the next fifth and the remainder separates it the same way; for CYP2D6 the red band runs at three times the library rate and the green band at half of it. Panels B and C carry the curves and the band rates for every endpoint.

The enrichment available is set by how rare the outcome is. Where half the library already carries the endpoint the score has little to sort: hepatotoxicity moves from 50 in 100 to 84 in 100, and cardiotoxicity at 30 micromolar from 82 to 92. The method earns its place on the rare outcomes, which are also the ones a triage step is run to find.

### 3.8 What everyone gains by contributing

The consortium results rest on the size of the pooled reference set, and the return on enlarging it is measured in Figure 1C. Growing the set from 127 to 5,953 compounds carrying a measured LD50 raises the share with an evaluable prediction from 3.3% to 46.5%, while accuracy rises from 85.4% to 90.9% over the first fivefold increase and then holds near 92%. A contributor gets that back immediately, because the method has no training step and a compound added to the pool is available to every query the next time one is run. The gain is shared and it is compounding: each contribution widens the region of chemistry in which everyone, the contributor included, receives an answer rather than a decline.

Toxicity data on discontinued compounds may be a practical place to start. When a compound shows toxicity the usual consequence is that it is deprioritized or its program is terminated, so the chemistry is less likely to be advanced. Contributors would still have to decide which records they are able to release, and the record below shows that decision has been the binding constraint rather than the technology. A finding that cost one company a program can still stop three others repeating it.

One caveat belongs on the record, and it bears on live chemistry rather than on a toxicity finding. Exchanging fingerprints in place of structures was once treated as sufficient protection on its own, and that position no longer holds. Trained on public data alone, and given no candidate library to match against, a decoder recovers 11.3% of one company’s proprietary compounds as exact structures from folded binary 1,024-bit fingerprints^24^. A later sequence model reports about 93% on compounds resembling its training set^25^. An organization contributing a series it is still working on should know that. For a compound it has stopped working on, which is what fills a toxicity pool, there is less at stake, although each contributor still has to decide what it can release.

The four counts described below disclose fewer features than a fingerprint. It is four small integers computed locally against the public index: families reached, proteins reached, matches, and how many of a 47-protein safety pharmacology antitarget list, distinct from the 19-protein comparator panel in section 2.4, those proteins include. They are not structural descriptors, and we are aware of no published method that recovers a structure from them, which makes them a lower-information contribution format rather than a demonstrated guarantee. On their own they reach a mean area under the ROC curve of 0.66 across the 28 endpoints, and 0.74 once coarsened heavy atom count, cLogP and molecular weight are added, against 0.72 for those size descriptors alone. A contributor who releases four integers in place of a structure therefore keeps about half of what the whole reverse screen is worth over molecular size, and releases nothing any published method inverts. The fingerprint remains the only format carrying both signals.

This has been attempted repeatedly and the binding constraint has not been technical. eTOX pooled 8,196 repeated-dose studies on 1,947 compounds from thirteen companies^26,33^. Of the 1,403 substances in its 2014 release, 515, or 37%, were held as confidential and carried no structure^26^. Its own methodologists then built a descriptor permutation that survived code-breaking challenges inside the consortium and preserved model quality. They reported that it could not be deployed, because the external audits the member companies would have required were unaffordable^27^. Its successor reported that 91% of donated data could not be shared even within the consortium^28^. Both projects released modeling software openly; neither released the pooled data, and the public eTOXsys instance has since been discontinued while the eTRANSAFE site no longer resolves^27,28^. The specific precedent for the present work is sharper still: eighteen European companies pooled acute toxicity data on 74 compounds in 2008. They used it to argue, with regulators from the US, Japan and Europe agreeing, that acute toxicity studies are not needed before first-in-human trials, and never released the compound-level data^29^. There is still no public cross-company acute toxicity database for pharmaceuticals.

The route the largest collaboration of this kind took was neither. MELLODDY trained across ten companies without descriptors leaving any owner’s infrastructure, and permuted fingerprint bits under a secret key^30^; eTOX withheld structures on its confidential tier and placed the data with a trusted third party^26^. Both are heavier machinery than a shared file. The measurements here describe what the lighter option costs.

### 3.9 The same rise on compounds from outside

Everything above was measured on compounds this work had access to. A separate group has since measured CYP3A4 inhibition on compounds it did not. What follows checks the association between family count and that one endpoint; it does not test the combined score of section 3.7 or the consortium predictor.

The OpenADMET consortium, a public effort that measures absorption and toxicity properties on common platforms and releases them openly,^31^ released CYP inhibition measurements from a single platform, of which 2,335 compounds carry a direct CYP3A4 pIC50^32^. A compound was counted as an inhibitor at pIC50 5.00 or above, equivalent to an IC50 of 10 uM or below, and that threshold was fixed before any compound was screened. On that rule 481 of the 2,335 are inhibitors, a rate of 20.6%.

Each compound was screened with the same fingerprint and the same procedure used throughout this work, and grouped by the number of protein families its matches reach. The index release used here is the 20 September 2026 release used everywhere else in this work. The percentage measured as a CYP3A4 inhibitor rises with that count (Table 1). Compounds reaching no family are inhibitors 7.3% of the time, about a third of the rate across the set, while those reaching sixteen families or more are inhibitors 24.6% of the time. Comparing families reached between inhibitors and the rest gives a Mann-Whitney p of 5.09e-22.

**Table 1.** Percentage of compounds measured as CYP3A4 inhibitors in the OpenADMET consortium dataset, by the number of protein families the reverse screen reaches. An inhibitor is a compound with a CYP3A4 pIC50 of 5.00 or above, a threshold fixed before screening. Index release 20 September 2026.

| Families reached | Compounds | Percent inhibitor |
| --- | --- | --- |
| 0 | 41 | 7.3 |
| 1 to 3 | 86 | 8.1 |
| 4 to 7 | 139 | 12.2 |
| 8 to 11 | 224 | 11.6 |
| 12 to 15 | 250 | 14.4 |
| 16 or more | 1,595 | 24.6 |
| All | 2,335 | 20.6 |

Read as a filter, a cut at sixteen families sets aside 740 compounds, just under a third of the set, at 12.03% inhibitor against 24.58% above the cut. The lower-reach group has a reduced inhibitor rate, although it still contains 89 of the 481 inhibitors.

Forty-three of the 2,335 are themselves ligands in the index the reverse screen searches, and 98 carry a measurement in the TOXRIC tables used above. Removing all 106 distinct compounds leaves the separation where it is: below a cut at sixteen families the inhibitor rate is 12.44 percent on 675 compounds against 24.45 percent on the 1,554 above it, with a Mann-Whitney p of 8.35e-20, against 12.03 percent and 24.58 percent on the full set.

### 3.10 A second retrieval arm

The index is searched by one fingerprint above. It can be searched by two. The same 27,910 ligands carry Morgan fingerprints of radius 2 over 2,048 bits, a different fingerprint from the radius 2 folded to 1,024 bits the consortium vote of section 2.1 uses, and a query can be screened against both, with the proteins the two sets of matched ligands were solved against combined and counted once each. The two similarities are on different scales, so one cutoff applied to both would mean two different things: the Morgan cutoff is set at 0.25, the value at which the Morgan arm admits the same median number of indexed ligands per compound as the pharmacophore arm admits at 0.50.

The first thing the second arm buys is coverage. 6,153 compounds reach no protein family through the pharmacophore arm, and 4,166 of them, 12.2% of every compound measured, reach at least one once the Morgan arm is included, gaining a median of seven families. They are mostly small flexible esters and ketones that carry too few pharmacophoric features to clear the pharmacophore cutoff while sharing enough substructure with an indexed ligand to clear the Morgan one: their Morgan fingerprints carry a median of 21 set bits against 36 across all compounds measured. Reach rises from a median of 10 families on the pharmacophore arm to 16 over both.

The second is accuracy against cytochrome inhibition. Carried as separate features rather than combined into one number, and scored by five fold cross validation so that the extra features cannot pay for themselves, the mean area under the ROC curve across the five cytochromes rises from 0.67 for families reached on the pharmacophore arm to 0.71 for both arms and both readouts. Counting proteins as well as families accounts for 0.03 of that and the Morgan arm adds a further 0.01 on top of it. Every per compound figure quoted elsewhere in this work is the pharmacophore arm at 0.50, which is what every published figure is measured on.

The arms have to be carried separately. Replacing the family count with the count over the combined arms loses, 0.64 against 0.67, and carrying the combined count with its protein count reaches 0.68, still below the 0.71 the four separate features reach. What the second arm adds is a second view, not a larger number.

## 4. Discussion

The method holds twenty-three years on. Rebuilt with an open fingerprint it reproduces the behavior reported then, including the property that makes pooling worthwhile: a larger shared reference set raises the share of compounds that can be answered for while leaving accuracy where it was. That the same machinery transfers without retuning to 28 endpoints it was never designed for says the premise holds, not that the implementation is clever. Like behaves like, over hepatotoxicity and eye corrosion and mutagenicity as much as over lethality.

The new quantity is target breadth computed from structure. Polypharmacology would lead one to expect that it predicts toxicity^8,9^. The useful part is that it is computed with no access to how much a compound has been tested, which is a claim measured promiscuity cannot make. That the relationship survives deleting the compound’s own scaffold, and deleting everything above 0.70 similarity to it, means it is not a lookup of close relatives. That the reverse screen adds to heavy atom count, molecular weight and cLogP on all 28 endpoints means it is not the physical property rule^10,11^ under a different name. The quantity carrying that gain is the count of distinct proteins reached; family count contributes little once the other retrieval measures are in the model, and it is kept because it is interpretable. The association between family count and CYP3A4 inhibition holds on 2,335 compounds measured by a separate consortium, none of which this work had seen. That check covers the family count alone, not the combined score and not the consortium predictor.

The endpoints that run backwards fit the same picture. Corrosion, irritation and mutagenicity are hazards of small reactive molecules, and compounds reaching no protein family are small. The count reports how many kinds of protein site a compound’s chemistry resembles; where the hazard does not come from binding at all, the trend inverts rather than disappearing.

Retrieval and the comparator together are more than either alone. A compound reaching many protein families, which a cross-family comparator also ranks highly at several safety panel proteins, inhibits CYP3A4 61.3% of the time, against 31.7% across the compounds that carry both signals. Requiring the retrieval and the comparator to agree raises confidence. On cardiotoxicity, double confirmation beats either signal alone, and the cardiac ion channels are in the panel the comparator was asked about.

Where this belongs in a program is early. ADME and toxicity are usually assessed once a series has been chosen and chemistry has been spent on it, which is the point at which a liability is most expensive to discover. Both signals here are computed from a two-dimensional structure, with no docking and no conformer generation at query time, so they run over a virtual library before any of it is made. Enrichment is largest exactly where it is most valuable to be early: at the androgen receptor ligand binding domain the score concentrates actives sixfold in the worst scoring tenth. Rare liabilities are the ones a program does not find by accident, and they are the ones that end it late. While the same score moves CYP3A4 inhibition from a third of the library to four fifths of the flagged tenth, the leverage there is smaller because a third of the library already inhibits; a filter cannot enrich what is already everywhere. We therefore position this as early triage over unmade chemistry rather than as a gate in front of an assay.

## 5. Conclusion

A compound’s toxicity can be gauged from the toxicities of the compounds it resembles; that was true in 2003 and it is true now on public data, over 28 endpoints the original work never touched. What is new is that a second question can be asked of the same compound at the same cost. How many different protein families does its chemistry reach in the crystallographic record? Compounds that reach widely are several times more likely to be active on cytochrome inhibition, on nuclear receptor binding and on clinical toxicity. The effect is specific to those endpoints; the number of distinct toxicity categories a compound is positive in does not rise with reach. That relationship holds after controlling for molecular size and lipophilicity, and it holds after deleting the compound’s own chemical series from the index. Used as a post-filter over a virtual library, the triage score concentrates the outcomes worth avoiding by three to six fold in the tenth it flags. That score combines retrieval measures, cross-family target preferences and physicochemical descriptors; consortium predictions are evaluated separately. Both halves are cheap, the consortium half improves as the shared reference set grows, the cross-family half is robust to how much of the present index is retained, and neither needs a protein structure or a docked pose.

The reference set is the model, and it grows only if people contribute to it. Twenty-three years ago we proposed that organizations pool encoded structure-property pairs and offered to host the pool, and the pool was never built. We renew that proposal here, with a working implementation and measurements of how reference set size affects coverage. Toxicity data on discontinued compounds may be the practical place to start: a compound that shows toxicity is usually deprioritized or terminated, so the chemistry is less likely to be advanced, although each contributor still has to decide which records it can release. A contributor can publish a fingerprint with its endpoints, or, for chemistry still in play, four integers that carry the cross-family signal and no structure at all. The software that reads this index also writes to it, at github.com/smuskal/toxpred, and interactive predictions and a route to contribute data are at toxpred.ai. We will host an open repository for contributed acute toxicity and endpoint data, accepting either form and not asking who contributed it, if there is interest from the organizations holding that data. Toxicology is the one place in drug discovery where a competitor’s failure is worth as much as one’s own.

## 6. Data, Code, and Availability

The acute toxicity reference set is the CATMoS rat oral collection^13^, distributed with the OPERA toolset^14^ and the further endpoints with TOXRIC^15^; both are public. The retrieval index is described in the reverse screening report^17^ and the cross-family comparator is available at familyfoundationmodel.com^22^.

The method is released as an open package, **toxpred**, at https://github.com/smuskal/toxpred. It fetches the public data when first run, scores a molecule or a file of molecules against the reference set, reports the further endpoints, and computes cross-family reach. Installation and use are documented there, and the figures in this manuscript are reproduced in that repository. The same method runs interactively at toxpred.ai, which scores a drawn or pasted molecule against the reference set and accepts a contributed record as structures, as fingerprints, or as the reach counts described in section 3.8.

For exact reproduction, two representations are used and they are not interchangeable. The consortium method of section 2.1 uses Morgan fingerprints of radius 2 folded to 1,024 bits. The reverse screen uses pharmacophore fingerprints from the published PharmCast release pharmcast_scp_v10.pt, for queries and for the index alike. Every retrieval used the Reverse Screen index release of 20 September 2026, which carries 27,910 ligands, the external set in section 3.9 included. Every indexed ligand at or above the 0.50 cutoff is kept, with no cap on the number of matches. Fingerprints are computed under rdkit 2025.9.6, the version both served implementations run; one of the descriptors the PharmCast surrogate reads differs under older rdkit releases. The results reported here were recomputed in that pinned environment on 13 October 2026. The site runs the same code against the current weekly release, which it names on every page. The package downloads both and verifies each against the checksums its publisher lists. Analysis code and every measured number reported here are held with the project record.

## 7. Acknowledgments

We thank Emerson Hsieh for discussions related to this data, and the OpenADMET consortium and Expansion Therapeutics for releasing it openly.

## 8. Competing Interests

S.M.M. is an employee of Eidogen-Sertanty, Inc. The remaining authors declare no competing interests.

## Notes

https://toxpred.ai/

https://github.com/NIEHS/OPERA

https://toxric.bioinforai.tech/home

https://reversescreen.ai/download.html

https://pharmcast.ai/

https://familyfoundationmodel.com/

### Competing Interest Statement

The authors have declared no competing interest.

### Summary of Updates

Reverse screen retrieval is no longer capped at 500 matches per query, and all cross-family figures were recomputed; the headline CYP3A4 band now runs from 2.5% to 51.2% rather than to 51.0%. Section 3.10 is new and adds a second retrieval arm over Morgan fingerprints, unioned with the pharmacophore arm. The external CYP3A4 check in section 3.9 was recomputed uncapped and on the same index release as the rest of the work, strengthening the separation. Supporting Information updated. Consortium results and Table S1 are unchanged.

