## Supporting Information: Figures S1 and S2, Tables S1 and S2 for "Toxicity by Consortium Revisited: Structural Neighbors and Cross-Family Reach"

### Supporting Information for "Toxicity by Consortium Revisited: Structural Neighbors and Cross-Family Reach"

Regenerated 14 October 2026 directly from data/curated/toxric_matrix.csv and data/curated/families_hit_full.csv, with the single blank-InChIKey row excluded. Analyzed data: 28 endpoints, 149,224 measurements, 34,162 compounds. At most two decimal places throughout.

#### Figure S1

Figure S1 puts twelve compounds from the top band of Figure 3 on the page. Each is a row of the TOXRIC tables, named in ChEMBL 37 and carrying the InChIKey it was selected by, and each reaches sixteen or more protein families. Felodipine is measured active on all three of the endpoints it was tested on, emodin on six of seven, and nitazoxanide on three of four. The denominator matters and is printed for every compound, because a compound measured on three endpoints and toxic on three is not the same claim as one measured on nine.

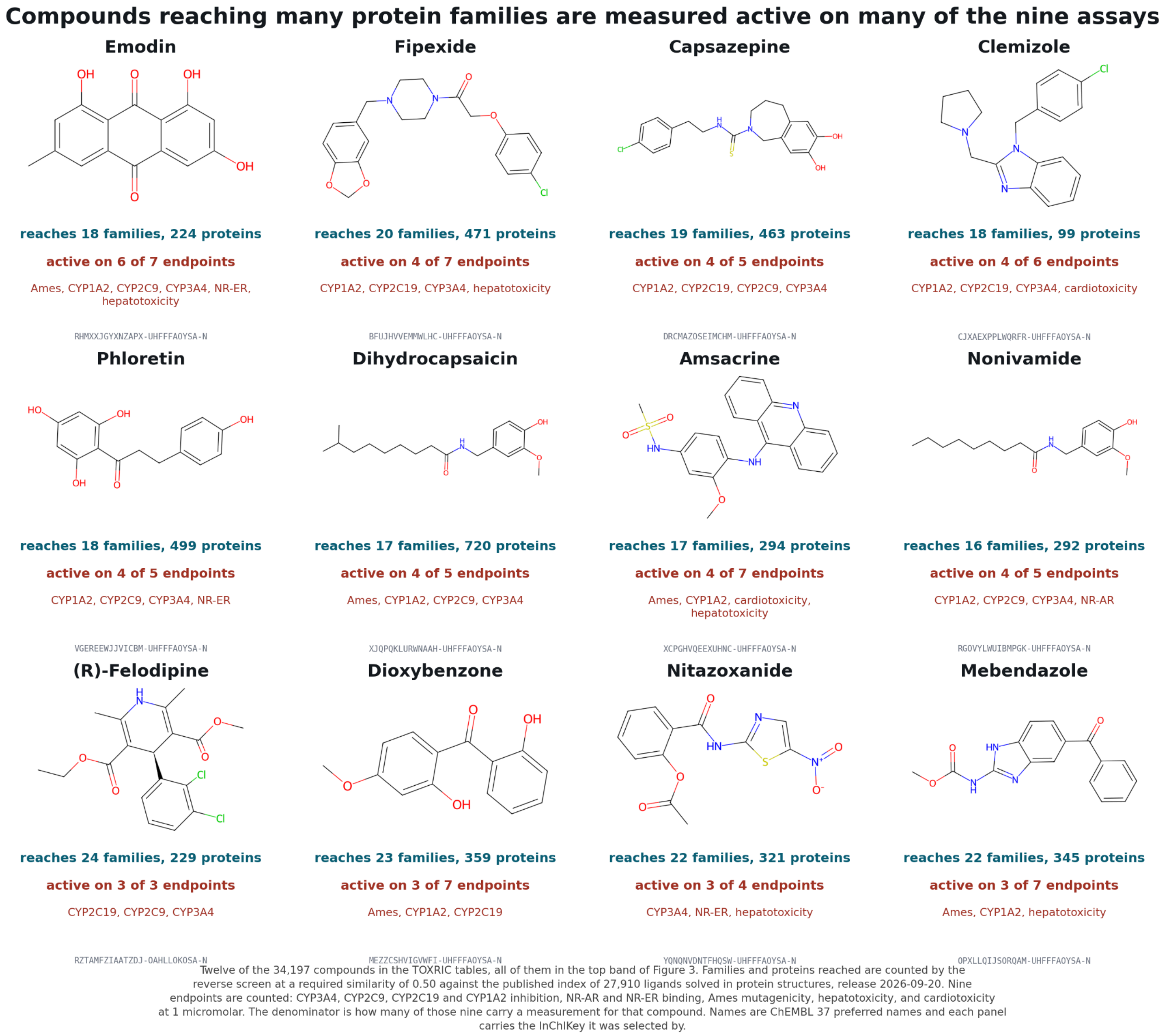


**Figure S1. Selected compounds with broad predicted reach and multiple positive assay results.** Twelve of the 34,197 compounds in the TOXRIC tables, every one of them in the top band of Figure 3. Families and proteins reached are counted by the reverse screen at a required similarity of 0.50 against the published index of 27,910 ligands solved in protein structures, release 20 September 2026. Nine endpoints are counted: CYP3A4, CYP2C9, CYP2C19 and CYP1A2 inhibition, NR-AR and NR-ER binding, Ames mutagenicity, hepatotoxicity, and cardiotoxicity at 1 micromolar. The denominator beside each compound is how many of those nine carry a measurement for it, so a compound measured on three endpoints is not read as though it had been measured on nine. Names are ChEMBL 37 preferred names and each panel carries the InChIKey it was selected by, so the same rows can be pulled from the same public tables. The panel is an illustration of the top band and carries no statistic of its own; the separation is measured in Figure 3 over all 34,162 compounds.

### Figure S2

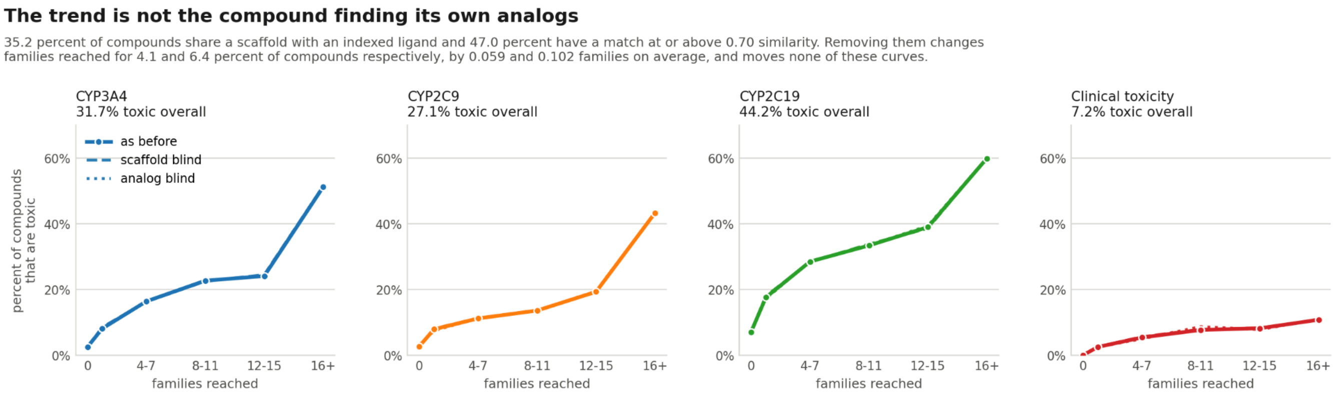


**Figure S2. The trend** is not a compound finding its own analogs. Each compound is screened three times: with the index as it stands, with every indexed ligand sharing the query's Bemis-Murcko scaffold removed first, and with every indexed ligand at or above 0.70 similarity to the query removed first. 35.2% of compounds share a scaffold with an indexed ligand and 47.0% have a match at or above 0.70, so there is a great deal to remove; removing it changes families reached for 4.1% and 6.4% of compounds respectively, by 0.06 and 0.10 families on average, and produces a largest single-band shift of 3.2 percentage points across all 28 endpoints under either blinding. For CYP3A4 no band moves by more than 0.4 of a percentage point.

### Table S1. Operating point for each of the 28 endpoints

Each endpoint uses the strictest required similarity that still answers for at least one compound in five; the value therefore differs by endpoint and is given here. Caught is the share of measured actives called active, and passed is the share of measured inactives called inactive, among the compounds answered for.

| Endpoint | Category | Required similarity | Answered (%) | Correct (%) | Actives caught (%) | Inactives passed (%) | Matthews correlation | Measured actives |
|---|---|---|---|---|---|---|---|---|
| Eye Corrosion | Irritation and Corrosion | 0.65 | 23.4 | 100.0 | 100.0 | 100.0 | 1.00 | 62 |
| Eye Irritation | Irritation and Corrosion | 0.65 | 26.8 | 98.6 | 99.2 | 96.5 | 0.96 | 260 |
| CYP1A2 | CYP450 | 0.75 | 21.9 | 96.1 | 90.7 | 98.6 | 0.91 | 161 |
| Ames Mutagenicity | Mutagenicity | 0.80 | 21.1 | 93.3 | 94.3 | 91.8 | 0.86 | 212 |
| CYP2C9 | CYP450 | 0.75 | 21.6 | 93.0 | 84.0 | 95.7 | 0.80 | 119 |
| CYP2C19 | CYP450 | 0.75 | 22.5 | 89.8 | 82.1 | 94.2 | 0.78 | 207 |
| CYP3A4 | CYP450 | 0.75 | 20.5 | 87.7 | 85.4 | 89.7 | 0.75 | 260 |
| NR-AR-LBD | Endocrine Disruption | 0.70 | 26.0 | 97.0 | 77.8 | 98.2 | 0.75 | 27 |

| | | | | | | | | |
|---|---|---|---|---|---|---|---|---|
| CYP2D6 | CYP450 | 0.75 | 21.2 | 92.8 | 68.8 | 97.0 | 0.70 | 80 |
| SR-MMP | Endocrine Disruption | 0.70 | 23.8 | 89.7 | 76.4 | 92.4 | 0.65 | 55 |
| NR-AhR | Endocrine Disruption | 0.70 | 23.9 | 94.7 | 60.6 | 98.0 | 0.64 | 33 |
| Respiratory Toxicity | Respiratory Toxicity | 0.75 | 26.6 | 85.3 | 87.5 | 79.0 | 0.63 | 56 |
| SR-ATAD5 | Endocrine Disruption | 0.70 | 25.1 | 97.7 | 50.0 | 99.5 | 0.62 | 16 |
| Hepatotoxicity | Hepatotoxicity | 0.75 | 22.8 | 80.9 | 83.1 | 78.8 | 0.62 | 77 |
| NR-AR | Endocrine Disruption | 0.70 | 23.5 | 94.9 | 48.5 | 99.0 | 0.60 | 33 |
| Cardiotoxicity-5 | Cardiotoxicity | 0.80 | 26.5 | 79.3 | 69.4 | 87.0 | 0.58 | 36 |
| Cardiotoxicity-10 | Cardiotoxicity | 0.80 | 26.5 | 79.3 | 79.6 | 78.8 | 0.58 | 49 |
| Cardiotoxicity-1 | Cardiotoxicity | 0.80 | 26.5 | 85.4 | 50.0 | 96.8 | 0.57 | 20 |
| NR-aromatase | Endocrine Disruption | 0.70 | 22.1 | 95.5 | 58.8 | 97.6 | 0.56 | 17 |
| SR-p53 | Endocrine Disruption | 0.70 | 22.8 | 96.0 | 47.6 | 98.9 | 0.56 | 21 |
| Carcinogenicity | Carcinogenicity | 0.65 | 21.2 | 75.5 | 78.6 | 71.4 | 0.50 | 28 |
| NR-ER-LBD | Endocrine Disruption | 0.70 | 25.0 | 95.3 | 43.5 | 98.2 | 0.48 | 23 |
| SR-ARE | Endocrine Disruption | 0.70 | 22.5 | 89.5 | 54.3 | 93.9 | 0.48 | 35 |
| NR-PPAR-gamma | Endocrine Disruption | 0.70 | 25.2 | 97.7 | 60.0 | 98.2 | 0.41 | 5 |
| Clinical toxicity | Clinical Toxicity | 0.70 | 20.8 | 91.8 | 40.0 | 96.4 | 0.40 | 5 |
| Cardiotoxicity-30 | Cardiotoxicity | 0.80 | 26.5 | 76.8 | 83.9 | 55.0 | 0.38 | 62 |
| NR-ER | Endocrine Disruption | 0.70 | 23.8 | 84.1 | 36.8 | 93.2 | 0.35 | 57 |
| SR-HSE | Endocrine Disruption | 0.70 | 22.0 | 95.9 | 36.4 | 97.9 | 0.34 | 11 |

Source: results/endpoint_operating_points.csv

### Table S2. Percent of compounds measured active, by protein families reached

Bands are the number of distinct level-2 protein families the reverse screen reaches at a required similarity of 0.50. Each cell gives the number of compounds in the band and the percent of them measured active.

| Endpoint | Compounds | Percent active | 0 | 1 to 3 | 4 to 7 | 8 to 11 | 12 to 15 | 16 or more |
|---|---|---|---|---|---|---|---|---|

| | | | | | | | | |
|---|---|---|---|---|---|---|---|---|
| Ames Mutagenicity | 6875 | 55.1 | 2030, 52.3% | 1758, 56.5% | 833, 66.3% | 655, 58.6% | 541, 53.8% | 1058, 48.0% |
| CYP1A2 | 9212 | 38.2 | 670, 10.9% | 859, 28.4% | 947, 39.7% | 965, 42.0% | 1020, 41.2% | 4751, 42.1% |
| CYP2C19 | 10095 | 44.2 | 718, 7.1% | 914, 17.6% | 984, 28.5% | 1040, 33.4% | 1119, 39.0% | 5320, 59.9% |
| CYP2C9 | 9405 | 27.1 | 750, 2.7% | 968, 7.9% | 986, 11.2% | 1021, 13.6% | 1044, 19.3% | 4636, 43.3% |
| CYP2D6 | 10201 | 13.4 | 740, 5.9% | 932, 10.7% | 983, 16.5% | 998, 15.7% | 1161, 14.1% | 5387, 13.7% |
| CYP3A4 | 10887 | 31.7 | 1212, 2.5% | 1338, 8.1% | 1036, 16.4% | 1067, 22.7% | 1066, 24.1% | 5168, 51.2% |
| Carcinogenicity | 1000 | 50.0 | 342, 52.9% | 231, 55.4% | 114, 50.0% | 85, 49.4% | 75, 46.7% | 153, 37.3% |
| Cardiotoxicity-1 | 1295 | 21.3 | 37, 5.4% | 89, 19.1% | 142, 16.2% | 171, 21.1% | 172, 12.2% | 684, 25.9% |
| Cardiotoxicity-10 | 1295 | 60.8 | 37, 37.8% | 89, 48.3% | 142, 57.0% | 171, 62.6% | 172, 64.5% | 684, 63.0% |
| Cardiotoxicity-30 | 1295 | 82.9 | 37, 62.2% | 89, 67.4% | 142, 83.8% | 171, 86.5% | 172, 87.8% | 684, 83.8% |
| Cardiotoxicity-5 | 1295 | 45.6 | 37, 21.6% | 89, 31.5% | 142, 40.1% | 171, 46.8% | 172, 42.4% | 684, 50.3% |
| Clinical toxicity | 1242 | 7.2 | 151, 0.0% | 160, 2.5% | 111, 5.4% | 142, 7.7% | 170, 8.2% | 508, 10.8% |
| Eye Corrosion | 2200 | 38.9 | 708, 82.2% | 379, 61.2% | 179, 11.7% | 209, 3.3% | 194, 1.5% | 531, 1.9% |
| Eye Irritation | 5052 | 74.7 | 2012, 94.9% | 1445, 90.7% | 397, 61.5% | 275, 28.7% | 267, 31.5% | 656, 22.4% |
| Hepatotoxicity | 2805 | 51.4 | 328, 47.3% | 314, 49.4% | 299, 62.2% | 328, 61.0% | 345, 53.9% | 1191, 47.1% |
| NR-AR | 6873 | 4.3 | 1912, 1.4% | 1569, 2.1% | 702, 4.6% | 613, 10.6% | 547, 5.7% | 1530, 7.2% |
| NR-AR-LBD | 6410 | 3.5 | 1863, 1.2% | 1491, 1.5% | 657, 3.3% | 576, 8.3% | 491, 5.3% | 1332, 6.1% |
| NR-AhR | 6200 | 11.8 | 1803, 7.5% | 1430, 12.9% | 623, 12.7% | 549, 13.1% | 487, 15.8% | 1308, 14.0% |
| NR-ER | 5875 | 12.9 | 1743, 7.2% | 1383, 13.0% | 616, 16.2% | 529, 19.7% | 458, 18.1% | 1146, 14.2% |
| NR-ER-LBD | 6589 | 4.9 | 1890, 2.2% | 1512, 4.2% | 665, 7.5% | 589, 10.4% | 527, 7.4% | 1406, 4.9% |
| NR-PPAR-gamma | 6106 | 2.6 | 1855, 1.3% | 1456, 1.9% | 624, 2.7% | 542, 2.0% | 462, 4.3% | 1167, 5.1% |
| NR-aromatase | 5515 | 5.2 | 1728, 2.0% | 1296, 3.9% | 537, 4.5% | 467, 5.4% | 389, 6.7% | 1098, 11.5% |
| Respiratory Toxicity | 1195 | 60.8 | 195, 71.3% | 185, 66.5% | 144, 57.6% | 168, 58.9% | 162, 54.9% | 341, 56.9% |
| SR-ARE | 5535 | 15.6 | 1755, 10.3% | 1350, 14.2% | 553, 15.7% | 443, 18.5% | 420, 21.2% | 1014, 23.3% |
| SR-ATAD5 | 6706 | 3.6 | 1918, 2.3% | 1548, 3.4% | 687, 4.4% | 607, 4.0% | 526, 5.7% | 1420, 4.1% |
| SR-HSE | 6138 | 5.4 | 1870, 3.8% | 1463, 5.2% | 617, 5.5% | 506, 5.5% | 473, 6.3% | 1209, 7.7% |
| SR-MMP | 5511 | 15.8 | 1685, 7.2% | 1323, 17.5% | 550, 16.7% | 468, 14.7% | 392, 20.2% | 1093, 25.4% |

| | | | | | | | | |
|---|---|---|---|---|---|---|---|---|
| SR-p53 | 6417 | 6.0 | 1884, 2.6% | 1482, 5.5% | 641, 5.5% | 567, 7.1% | 510, 11.0% | 1333, 9.5% |

Source: recomputed from data/curated/toxric_matrix.csv and data/curated/families_hit_full.csv, screened with no cap on the number of matches. Totals: 149,224 measurements over the 28 endpoints.
